# High-resolution mapping of RNA structural maturation during Cas9 assembly with ABEL-FRET

**DOI:** 10.64898/2026.09.25.754466

**Authors:** Bok-Eum Choi, Hugh Wilson, Quan Wang

## Abstract

The structural flexibility of RNA is essential for forming ribonucleoprotein (RNP) complexes, which regulate diverse biological processes. This intrinsic property permits RNA to act as a dynamic scaffold along the assembly pathway as it folds into a specific structure for initial recognition by protein and undergoes conformational rearrangements for functional maturation as a complex. Yet, RNA flexibility and RNP multicomponent assembly create significant obstacles for traditional structural methods. To overcome these challenges, we applied recently developed ABEL-FRET spectroscopy to measure tether-free single-molecule Förster resonance energy transfer (smFRET) over extended observation times. Furthermore, ABEL-FRET enables the unique ability for simultaneous measurements of ultrahigh resolution smFRET and hydrodynamic size of individual complexes, which offers distinct advantages for studying dynamic RNA molecules that undergo assembly via sequential binding events. Using ABEL-FRET, we explored how the guide RNA (gRNA) of CRISPR genome editing system folds and modulates its structural flexibility to carry out the roles required for each assembly state from its unbound *apo* form to the functional Cas9 RNP state for target DNA cleavage. Multi-perspective view of gRNA structure gained by probing its two primary functional domains enabled to capture dramatic changes in gRNA flexibility that are highly dependent on its specific structural domains as well as assembly states. Collectively, our work with ABEL-FRET highlights the intrinsic link between the structural flexibility of RNA and its functionality in RNP assembly.

## Introduction

Non-coding RNA performs essential functions for diverse biological processes, primarily as a ribonucleoprotein (RNP) by forming a complex with specific proteins^1,2^. Despite its functional importance, a comprehensive understanding of how RNA folds and assembles with proteins is still lacking due to limited structural information. This reflects challenges with traditional structural methods for studying biological systems that involve structural flexibility and compositional heterogeneity. Given this intrinsic complexity, single-molecule Förster resonance energy transfer (smFRET) has recently emerged as a new tool to study RNA structures and RNP assembly mechanisms^3–5^ as it reveals conformational dynamics and distributions. Although smFRET has provided insight that was not accessible with other methods, conventional smFRET measurements alone are often inefficient for multicomponent systems such as RNP assembly, which require correlated information on both conformational changes and binding events.

To address the limitations of current techniques for studying RNA and RNP assembly, we applied our ABEL-FRET that provides a tether-free platform for smFRET measurements with ultrahigh resolution and additional hydrodynamic profiles^6^. Our motivation was driven by its unique ability in performing simultaneous measurements for ultrahigh-resolution FRET efficiency (parameter for conformation and dynamics, *E*_FRET_), diffusion coefficient (parameter for size and shape, *D*), and electrokinetic mobility (parameter for surface charge density, *µ*) of single molecules in solution^7,8^. When the labeled biomolecule of interest forms a complex with the unlabeled assembling partners, not only the structure but also the diffusivity and electric-field-induced motion are likely changed. In this respect, our approach with ABLE-FRET, which provides multiple parameters, can effectively determine the conformational changes of the molecule with *E*_FRET_ while monitoring *D* and *µ* that are altered by assembly with binding partners^6^. Therefore, ABEL-FRET has the potential to visualize the architecture of RNA as it takes an assembly journey from the inactive precursor to the final functional RNP complex via sequential binding events. Furthermore, changing labeling positions of a donor-acceptor FRET dye pair within RNA permits us to gain a multi-perspective view of RNA structure along its assembly into RNP.

As an experimental RNP model for ABEL-FRET measurements, we chose the CRISPR/Cas9 genome editing system which works as a RNP complex with a Cas9 nuclease and a guide RNA (gRNA)^9,10^. Since Cas9 RNP is widely used in the scientific community, extensive efforts have been devoted for understanding its structure^11–14^ and underlying mechanisms using various methods including smFRET^15–19^. Although previous studies suggested the importance of gRNA structure in the Cas9 RNP assembly process^12–14^, none of these works have directly characterized its initial folding before binding to Cas9 and the subsequent conformational rearrangements upon the sequential assembly steps. In fact, our current understanding of gRNA is heavily reliant on its static crystal structure in the Cas9-bound state, which often leads to overlooking not only the structural integrity of gRNA in the unbound *apo* state but also intrinsic conformational dynamic and heterogeneity as RNA molecules.

Like other RNP complexes^20,21^, gRNA undergoes a sequential and hierarchical process for proper RNP assembly, which involves the initial RNA folding and subsequent rearrangements during assembly. First, two RNA molecules, CRISPR RNA (crRNA) and trans-activating crRNA (tracrRNA), fold together into a partial duplex gRNA (tracrRNA:crRNA), where sequence-conserved tracrRNA provides a ‘scaffold’ for the Cas9 binding, and crRNA carries the 5’ end 20-nucleotide (nt) ‘spacer’ sequence for target DNA recognition^9,22^. For simplicity, these two RNAs are often synthesized as a single gRNA (sgRNA) by linking crRNA and tracrRNA sequences with a tetra loop^22^. Next, this pre-folded gRNA assembles into an RNP complex with a Cas9 protein. Finally, the Cas9-gRNA RNP binds to the target DNA for double-stranded (ds) break.

In this study, we used our ABEL-FRET to investigate the local conformations adopted by scaffold and spacer, which are two primary functional regions of gRNA throughout the assembly process. For the scaffold region, we monitored the assembly pathway starting from tracrRNA that initially existed as a heterogeneous ensemble of interconverting states then transformed into a homogeneous stable structure as a part of gRNA scaffold. When the target-specific spacer region at the 5’ end sgRNA was probed, we observed that it exists as a highly flexible extension. This additional view in the *apo* gRNA envisions that gRNA initially folds into a hybrid structure of well-defined scaffold and flexible spacer at the 5’ end. Along with the gRNA assembly into Cas9 RNP, its rigid scaffold provides the docking site for Cas9 protein binding that induces narrowing of the conformational adoptions arising at the spacer sequence. The succeeding target DNA binding drives the spacer transition to the thermodynamically stable RNA-DNA heteroduplex that can induce further conformational changes in Cas9. Taken together, our work using ABEL-FRET visualizes the previously hidden gRNA folding and its rearrangements via sequential assembly. More importantly, our approach enables a move beyond static structural models to a functional landscape view as ABEL-FRET visualizes how gRNA modulates its flexibility along the assembly into RNP, which serves as an exciting addition to our understanding of gRNA structure–function relationship.

## Results

### ABEL-FRET monitors the initial gRNA folding and subsequent rearrangements along the sequential RNP assembly

For the demonstration of our approach using ABEL-FRET, we first probed the sequence-conserved scaffold using tracrRNA labeled at its 5’ end with Cy3 and sulfo-Cy5 (sCy5) dyes where crRNA:tracrRNA (gRNA) partial duplex forms (Fig. 1A). This labeled tracrRNA remained fully functional as a part of gRNA and Cas9 RNP that maintained the enzymatic activity for target DNA cleavage (Fig. S1). To capture the conformational transitions from the initial tracrRNA state to functional RNP complex, samples for ABEL-FRET measurements were prepared with the labeled tracrRNA by performing a step-by-step assembly process starting from tracrRNA only (tracRNA) into gRNA (tracrRNA + crRNA) and then RNP with dCas9 (tracrRNA + crRNA + dCas9) (see Methods for details). Due to the catalytic inability of dCas9^9,13^, the subsequent incubation with target dsDNA (tracrRNA + crRNA + dCas9 + DNA) resulted in the target-bound RNP complex formation without cleavage. These preassembled tracrRNA samples were then diluted to picomolar concentrations and then directly subjected to ABEL-FRET that provides simultaneous measurements for conformations with *E*_FRET_ and assembly states with *D* and *μ* as shown in the representative time trace (Fig. 1C).

**Figure 1.**
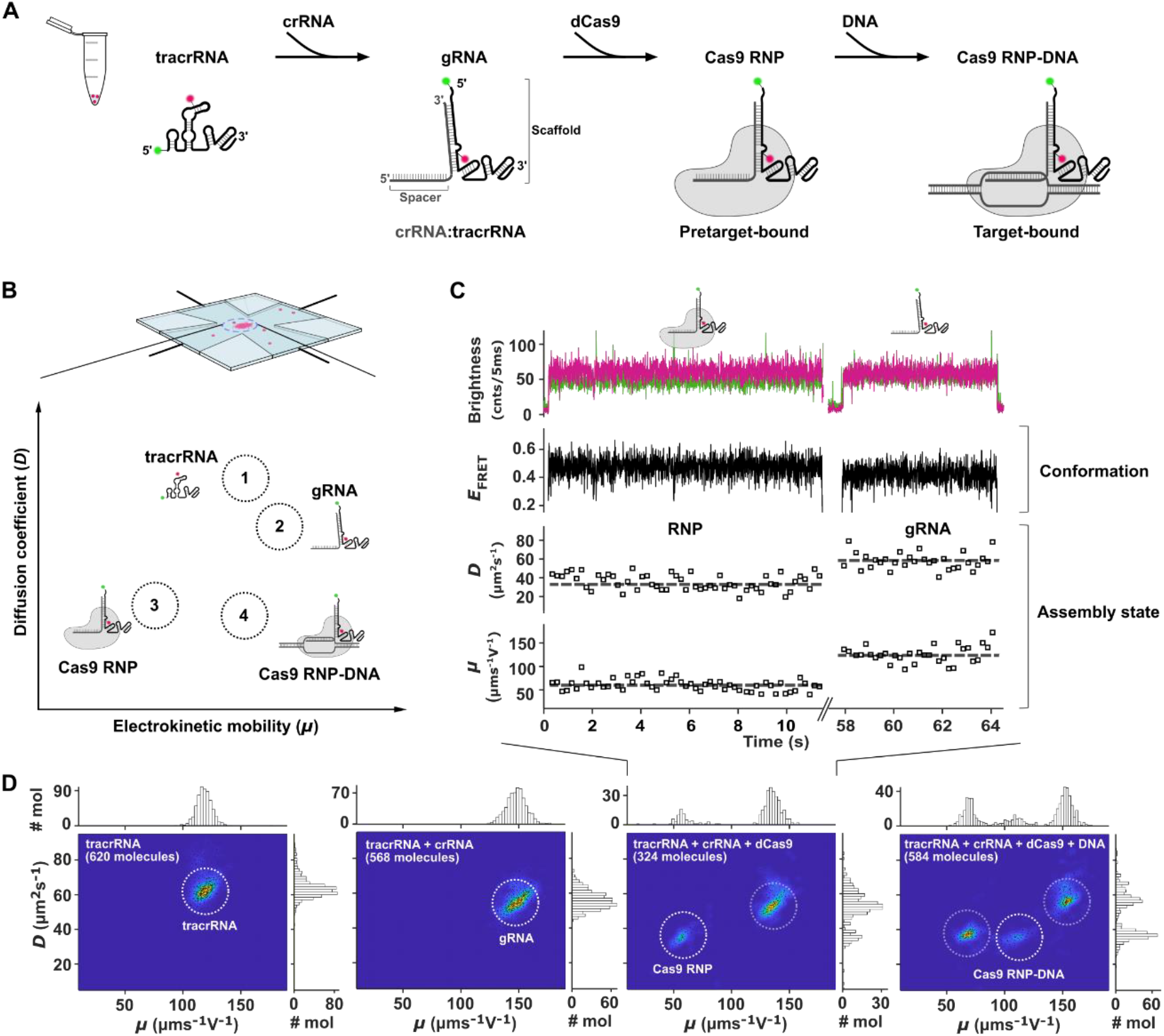
Experimental design and concept for conformational measurements of gRNA along its assembly pathway using ABEL-FRET. **(A)** CRISPR-Cas9 assembly steps from tracrRNA into gRNA (crRNA:tracrRNA), Cas9 RNP (gRNA-Cas9), and target-bound RNP. ABEL-FRET measurements were performed for tracrRNA before and after each addition with its assembling partners (crRNA, dCas9, and target dsDNA). **(B)** Expected diffusivity (*D)* and mobility (*μ)* tendency of tracrRNA throughout its assembly process. **(C)** A representative single-molecule time trace collected with ABEL-FRET for the tracrRNA sample assembled with crRNA and dCas9. Two molecules were sequentially trapped for measurements. Brightness of donor (green) and acceptor (red) channels are quantified by the number of photons per 5 ms time bins. *E*_FRET_ (10-ms bins), *D* (200-ms bins), and *μ* (200-ms bins) were simultaneously measured for single molecules. Horizontal dashed lines on *D* and *μ* traces indicate assembly states of trapped molecules, Cas9 RNP and gRNA, respectively. **(D)** Single molecule 2D *D*-*μ* scatter plots for the entire population (FRET and donor-only molecules) collected from tracrRNA samples pre-assembled by itself or with other assembly components. Density plots are overlaid with scatter plots, and marginal histograms along *D* and *μ* are shown. Both FRET and donor-only molecules are plotted here. The mean values of *D* and *μ* for each assembly state observed from all the samples are contained in Table S2.

As tracrRNA associates with it binding partners, it can have four distinct assembly states regarding the number of bound components from the initial state as tracrRNA itself to the final state as a part of target-bound Cas9 RNP (Fig. 1A). Since nucleic acids (DNA and RNA) with phosphate backbone are known to be highly negative molecules compared to proteins, an electric field in the trap was generated to capture negatively charged molecules including tracrRNA. As the assembly process proceeded, additional components were associated with tracrRNA and expected to increase the overall molecular size which is evident by the decrease in diffusivity (*D*) (Fig. 1B). The effect of binding with each assembling partner on electrokinetic response (*μ*), which is dependent on the surface charge density, is relatively ambiguous to predict. However, it is reasonable to anticipate the reduced *μ* as gRNA (tracrRNA:crRNA) assembled RNP with a Cas9 protein (Fig. 1B).

In fact, ABEL-FRET measurements for these preassembled tracrRNA samples were in good agreement with our prediction on *D* and *μ* trends shifted along the sequential assembly. In the *D*-*μ* two-dimensional (2D) plot, the sample that contained only tracrRNA exhibited one major group which then shifted towards the lower *D* and higher *μ* when crRNA was added (tracrRNA vs. tracrRNA + crRNA in Fig. 1D and Table S2). This indicates that all tracrRNA hybridized with crRNA and formed a partial duplex gRNA that has a larger size with a higher negative charge density on the molecular surface compared to tracrRNA. The pre-annealed gRNA was then incubated with dCas9 (tracrRNA + crRNA + dCas9), leading to the appearance of RNP population in addition to gRNA (Fig. 1D). As expected, the protein binding resulted in a decrease in both *D* and *μ* compared to gRNA (Table S2). Finally, when dsDNA target was added, we observed the target-bound RNP (RNP-DNA) as the third population which was apart from the free RNP population by its enhanced *μ* with a slightly decreased *D* (tracrRNA + crRNA + dCas9 + DNA in Fig. 1D and Table S2).

Under our experimental conditions, not all gRNA molecules bound to dCas9 and target DNA, which led to the coexistence of different assembly states. This compositional heterogeneity is a common feature when studying multicomponent biological systems and can be problematic for accurate characterization with conventional methods. In this regard, our results highlight that ABEL-FRET is advanced with its capability in resolving compositional heterogeneity by monitoring *D* and *μ* without any complication in addition to *E*_FRET_ measurements.

### The tracrRNA folds from a heterogeneous conformational ensemble into a homogeneous gRNA scaffold via hybridization with crRNA

To overview the distributions of tracrRNA conformations, we first plotted *E*_FRET_ histogram of tracrRNA molecules for each assembly state with a Gaussian fit (Fig. 2A). When there was no obvious sign of heterogeneity in regard to assembly states (tracrRNA & tracrRNA + crRNA in Fig. 1D), all of the molecules collected from each sample were determined to be in the same assembly state, therefore the entire population was plotted (tracrRNA & gRNA in Fig. 2A). First, the *E*_FRET_ histogram of tracrRNA exhibited multiple broad peaks with a major population around 0.82 and minor populations around 0.53 and 0.92 (Fig. 2A, first histogram from the left). Broad distributions in the histogram reflect dynamic transitions of 5’ end tracrRNA between different FRET states as shown in its real-time traces (Fig. 2B, left trace & Fig. S2). These observations imply that the 5’ end tracrRNA exists as a conformational ensemble that can interconvert. Strikingly, when this 5’ end tracrRNA hybridized with crRNA and formed a duplex within the gRNA structure, it gave rise to one major *E*_FRET_ population at 0.42 (Fig. 2A, second histogram). This RNA-RNA duplex appeared to be relatively static (Fig. 2B, right trace), indicating a conformationally constrained state as a part of gRNA scaffold.

**Figure 2.**
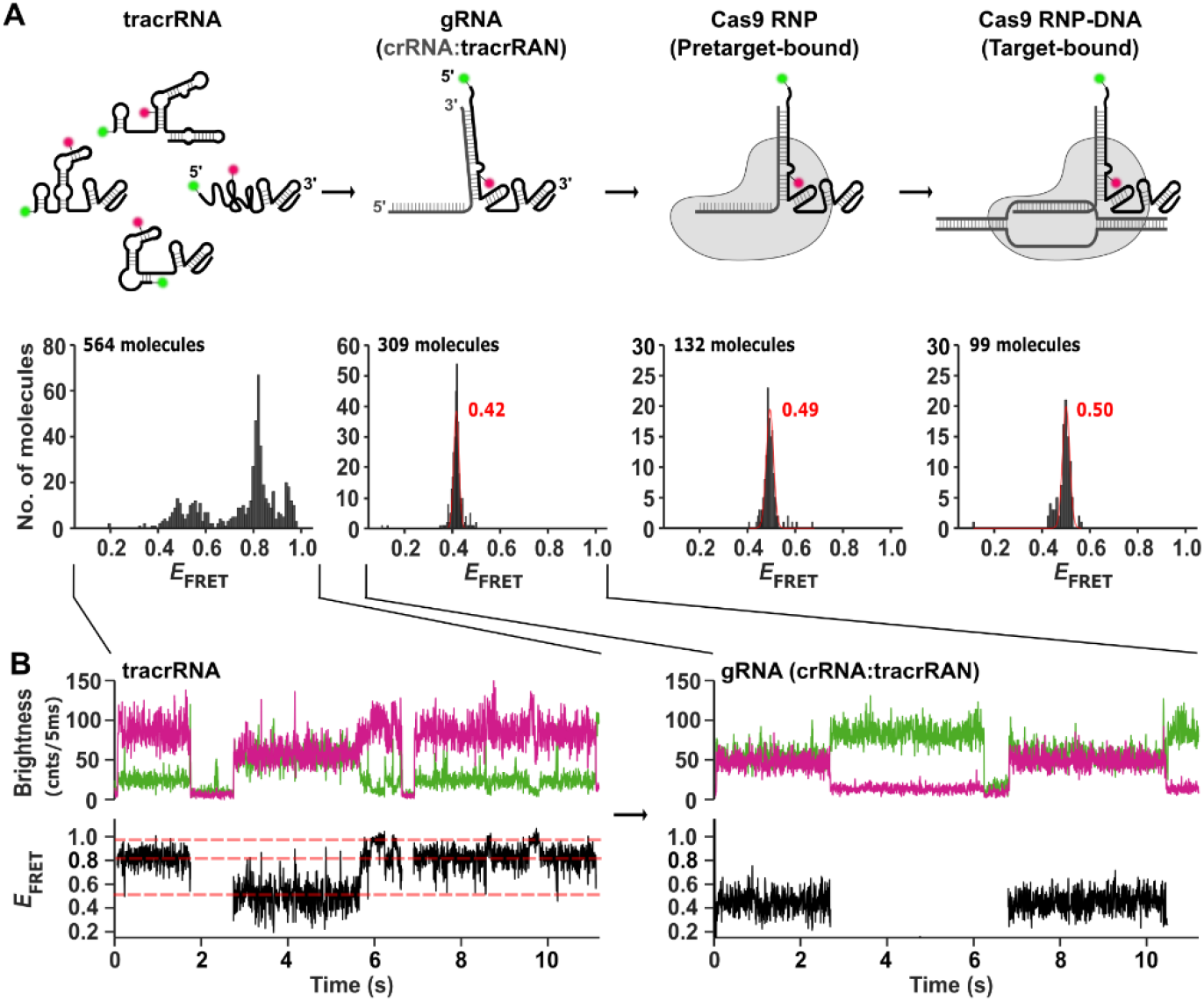
Conformational rearrangements of tracrRNA scaffold along its functional maturation into gRNA and Cas9 RNP. **(A)** Top: illustration of tracrRNA conformational rearrangements along the assembly process. The tracrRNA structures presented here were based on its predicted secondary structures by OligoAnalyzer available from the Integrated DNA Technologies (IDP) website. Bottom: *E*_FRET_ histograms of individual tracrRNA molecules and those in form of gRNA (crRNA:tracrRNA), RNP (gRNA-dCas9), and RNP-DNA. Histograms for RNP and RNP-DNA were generated with multiple data sets. Gaussian fits for gRNA, RNP, and RNP-DNA are shown in red with mean *E*_FRET_ values and standard deviations. Histogram for tracrRNA exhibits at least three *E*_FRET_ populations. **(B)** Examples of time traces collected for 5’ end tracrRNA before (left) and after hybridization with crRNA to form a partial duplex gRNA (right). Horizontal dashed lines in time trace for the single molecule of tracrRNA indicate multiple *E*_FRET_ states.

We observed the coexistence of multiple states after the protein addition (tracrRNA + crRNA + dCas9 & tracrRNA + crRNA + dCas9 in Fig. 1D), which accompanied with multiple FRET populations (Fig. S3). Based on their *D* and *µ* values, each FRET population was therefore assigned with the assembly state. The *E*_FRET_ histogram for tracrRNA molecules either in RNP or RNP-DNA state exhibited one major population (Fig. 2A). Upon the RNP formation with dCas9 (dCas9 RNP), we observed the *E*_FRET_ shift to 0.49, indicating that the gRNA partial duplex rearranged its conformation (third histogram in Fig. 2A). These results were further confirmed with Cas9 RNP that exhibited *E*_FRET_ at 0.49 for the partial duplex conformation (Fig. S4). To capture the target-bound state, gRNA was assembled with dCas9, followed by the incubation with target dsDNA. Our results suggest that the target binding had no significant effect on the conformation of gRNA partial duplex as shown with *E*_FRET_ at 0.50 (Fig. 2A, last histogram) which is very close to what we observed in the pretarget-bound state (0.49). Therefore, gRNA scaffold, particularly the partial duplex, is unlikely to play a major role in target recognition and binding. Taken together, our data highlights not only the unique ability of ABEL-FRET in simultaneous measurements for conformations and assembly states but also its ultrahigh resolving power in *E*_FRET_ measurements with extremely narrow distributions.

### Probing the target-specific spacer provides an additional angle to visualize gRNA with a flexible 5’ extension

We next located the Cy3-sCy5 dye pair at the gRNA spacer region where the 20-nt programmable sequence directs Cas9 to its complementary target DNA sequence for cleavage. In this case, we used sgRNA that required the assembly process with one less step by connecting the tracrRNA sequence to the crRNA sequence with the 5’ end 20-nt spacer (Fig. 3A). The fusion of two RNA molecules with minor sequence modifications and change in labeling locations are unlikely to induce a significant variation in gRNA hydrodynamic properties. Therefore, we anticipated the similarity between a partial duplex gRNA (crRNA:tracrRNA) and sgRNA in respect to their *D*-*μ* shifting patterns along the assembly process. Indeed, sgRNA also exhibited decreased *D* values as it assembled into RNP and target-bound RNP (RNP-DNA) which were distinguished by their *μ* values (Fig. 3B and Table S2).

**Figure 3.**
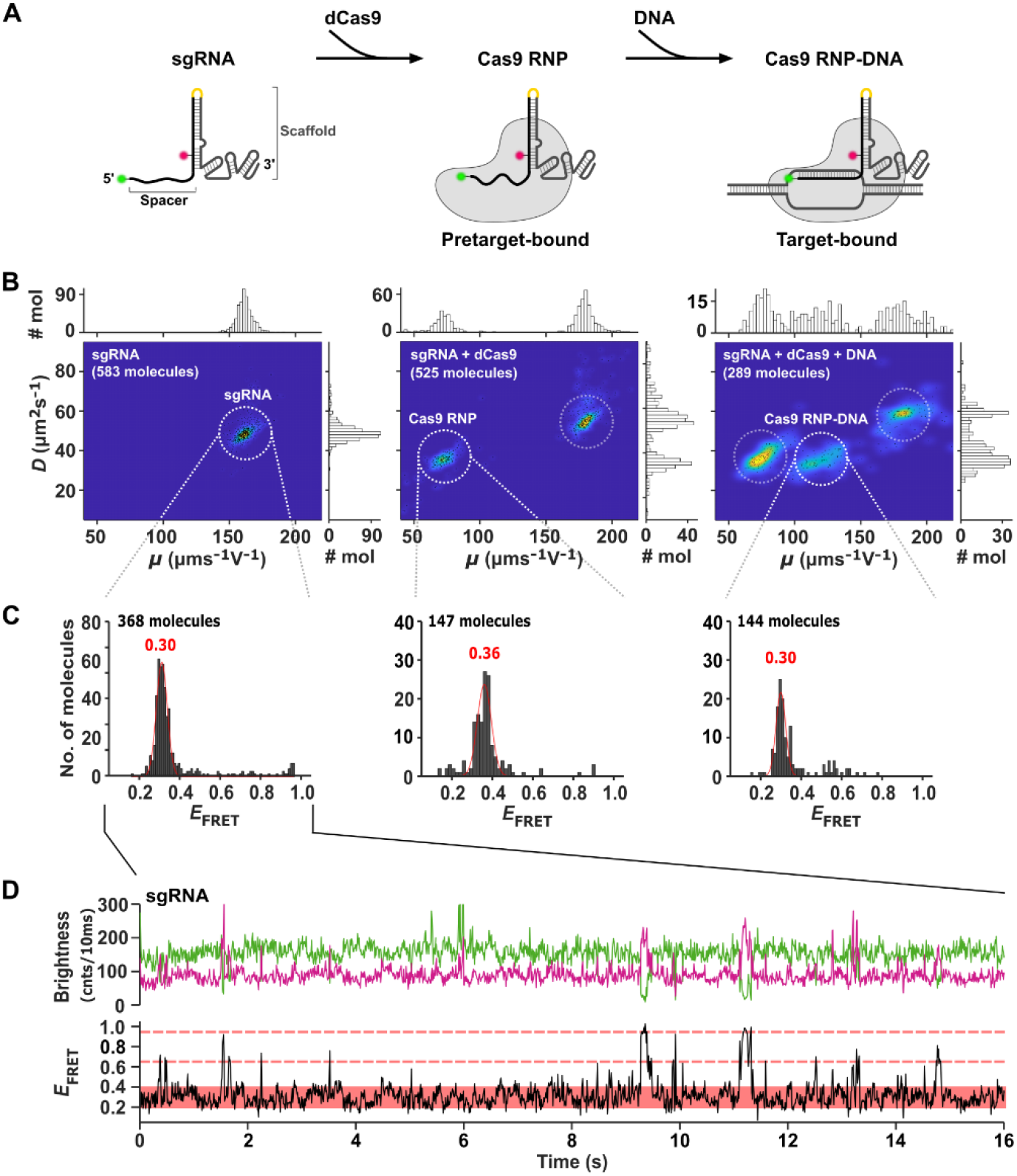
ABEL-FRET measurements of 5’ end sgRNA spacer throughout its assembly process. **(A)** Illustration of sgRNA assembly pathway. In sgRNA, crRNA and tracrRNA sequences were fused together by a tetra loop (yellow). **(B)** Single-molecule 2D *D*-*μ* density plots overlaid with scatter plots for the entire populations (FRET and donor-only molecules) from sgRNA samples before and after incubating with dCas9 and target DNA. Marginal histograms of *D* and *μ* are shown. The mean values of *D* and *μ* for each assembly state observed from all the samples are contained in Table S2. **(C)** *E*_FRET_ histograms of individual sgRNA molecules and those selected for RNP and RNP-DNA states accordingly with *D* and *μ* parameters. Gaussian fits are shown in red with mean *E*_FRET_ values and standard deviations. **(D)** Examples of time traces collected for 5’ end sgRNA in the *apo* state with continuous fluctuations in the low FRET state (red box) and occasional transitions to mid-high FRET states (red dashed lines). Quantified analysis for dynamic behaviors can be found in Fig. S5.

The histogram of 5’ end gRNA spacer in the initial state showed a major population at the low FRET state with a mean *E*_FRET_ at 0.30 (Fig. 3C, left histogram), which was given by the 5’ end fluctuating in the low FRET state with occasional transitions into mid-to-high FRET states (Fig. 3D and S5). When the RNP complex was formed, the mean *E*_FRET_ increased to 0.36 with dCas9 (Fig. 3C, middle histogram) or 0.37 with Cas9 (Fig. S6). On the other hand, the target dsDNA binding led to a decrease in the mean *E*_FRET_ at 0.30 (Fig. 3C, right histogram) which corresponded to the RNA-DNA heteroduplex formed between the 5’ end gRNA spacer and target DNA strand. This *E*_FRET_ value at 0.30 in the target-bound state is identical to what we observed for the 5’ end spacer in the initial state as a free gRNA, suggesting that the 5’ end gRNA tends to be extended even before Cas9 binding. However, its dynamic behaviors in the *apo* and target-bound states were very distinct from each other as discussed in the following section. Taken together, our measurements for the 5’ end gRNA in the *apo*, pretarget-bound, and target-bound states provide an additional view to feature gRNA as it initially folds into the structure that combines a highly stable double-stranded stem (scaffold region) with a flexible or dynamic single-stranded 5’ end segment (spacer region).

### ABEL-FRET visualizes that the flexibility of gRNA spacer is drastically reduced upon the target binding

Previous crystal structure of Cas9 RNP in the pretarget-bound states indicates that the 5’ end 20-nt gRNA spacer comprises as a disordered non-seed region (5′ 10-nt) in addition to a preordered A-form “seed” region where the initial interaction with its potential target occurs^11^. This disordered region, which was not visible in the structure, became the ordered A-form helical structure upon the target DNA binding (Fig. 4A). Our ABEL-FRET revealed that previously observed disordered property of pretarget-bound 5’ end gRNA is associated with conformational dynamics (Fig. 4B). As shown in a representative trace, the 5’ end sgRNA in the pretarget-bound state exhibited the anti-correlated fluctuations between donor and acceptor signals (top panel in Fig. 4B & S8), which was absent in the target-bound state (bottom panel in Fig. 4B). We further analyzed these time traces to extract quantitative parameters by fitting them into cross-correlation curves. Unlike the target-bound 5’ end without a clear indication for dynamics, the analysis revealed that the 5’ end gRNA in the pretarget-bound state experienced conformational dynamics on the millisecond timescales (Fig. 3C), which was the case for most sgRNA molecules (Fig. 4C).

**Figure 4.**
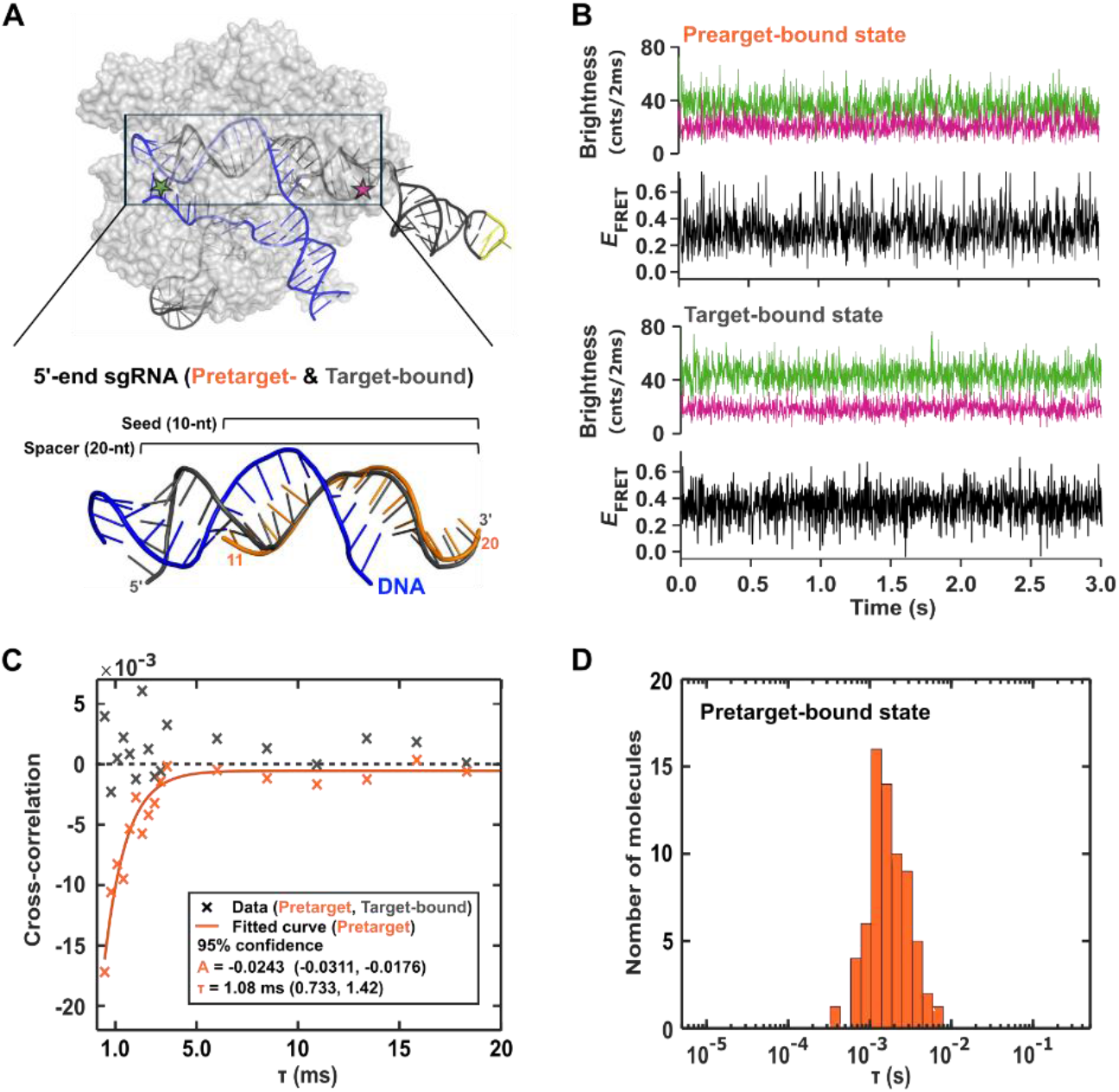
Dynamic behaviors of sgRNA spacer before and after target binding. **(A)** Top: Crystal structure of Cas9 RNP bound to target DNA (blue) (PDB: 5F9R) with the labeling position of sgRNA (black with a tetra loop in yellow). Bottom: Superimposed 5’ end sgRNA in the pretarget (orange, PDB: 4ZT0) and target-bound states (grey) that forms an RNA-DNA heteroduplex with the target DNA (blue). **(B)** Representative single-molecule time traces of sgRNA in RNPs before (top; pretarget-bound state) and after target binding (bottom; target-bound state). **(C)** Cross-correlation curve for the pretarget sgRNA (orange) with dynamic parameters presented in **B** compared to one in the target-bound state (gray) which was not fit to the curve. **(D)** Histogram of timescale for observed dynamics at the 5’ end sgRNA in the pretarget-bound state.

In the *apo* state, the 5’ end of gRNA appeared to be very dynamic as it continuously fluctuates in the low FRET (0.25-0.35) state with sudden jump to mid-to-high FRET (0.6-0.8) states (Fig. 3D and S5). This reflects its highly flexible structure, which explains why the *apo* gRNA structure is currently unavailable. Furthermore, our cross-correlation analysis indicates that the observed 5’ end fluctuation in the low FRET state occurs on hundreds of millisecond scales (Fig. S5), which is much slower than what we observed for the 5’ end spacer in the gRNA-Cas9 RNP complex (Fig. 4C). Eventually, the dynamic of gRNA was completely suppressed after the target DNA binding to the spacer. These results show that the 5’ end gRNA spacer shifts from a flexible to a more structured state along its assembly process. More importantly, our ABEL-FRET for real-time measurements revealed how the conformational dynamics at the 5’ end programmable spacer sequence changes, which were hidden by static snapshots in classical crystal structures.

## Discussion

Here, we demonstrated our ABEL-FRET as an advanced tool for studying RNA molecules that are involved in the assembly of RNP complexes. The unique ability of ABEL-FRET for simultaneously measuring multiple parameters (*E*_FRET_, *D*, and *μ*) of single molecules in real-time^6^ enabled direct characterizations of the flexible RNA structure with *E*_FRET_, while monitoring binding processes with *D* and *μ* changes. As an RNP model, gRNA-Cas9 complex served as an ideal system since extensive literature aided in interpreting our ABEL-FRET data while still requiring continuous research to fill knowledge gaps mainly driven by overlooking the dynamic nature of gRNA in its unbound *apo* as well as Cas9 RNP state. To address these knowledge gaps, we utilized our ABEL-FRET to examine the dynamic landscape of gRNA and its modulation in response to sequential bindings with Cas9 and target DNA as depicted in Fig. 5. In this work, we focused on two primary functional domains of gRNA that have been known to play critical roles in binding events, the conserved structural scaffold (tracrRNA) for Cas9 binding and the programmable spacer sequence (crRNA) for target DNA recognition^9,22^. Two independent data sets using gRNA molecules with different labeling positions exhibited consistency in shifting patterns of their hydrodynamic properties (*D* and *μ*) along the assembly with Cas9 (and dCas9) protein and target DNA (Fig. 1D & 3B), which validated our *D*-*μ* based assignments of assembly states for molecules. On the other hand, we observed distinct structural features (*E*_FRET_) for gRNA molecules with the same dye pair at different labeling positions, confirming that the observed FRET changes corresponded to structural rearrangement and not due to fluorescence dye artifacts.

**Figure 5.**
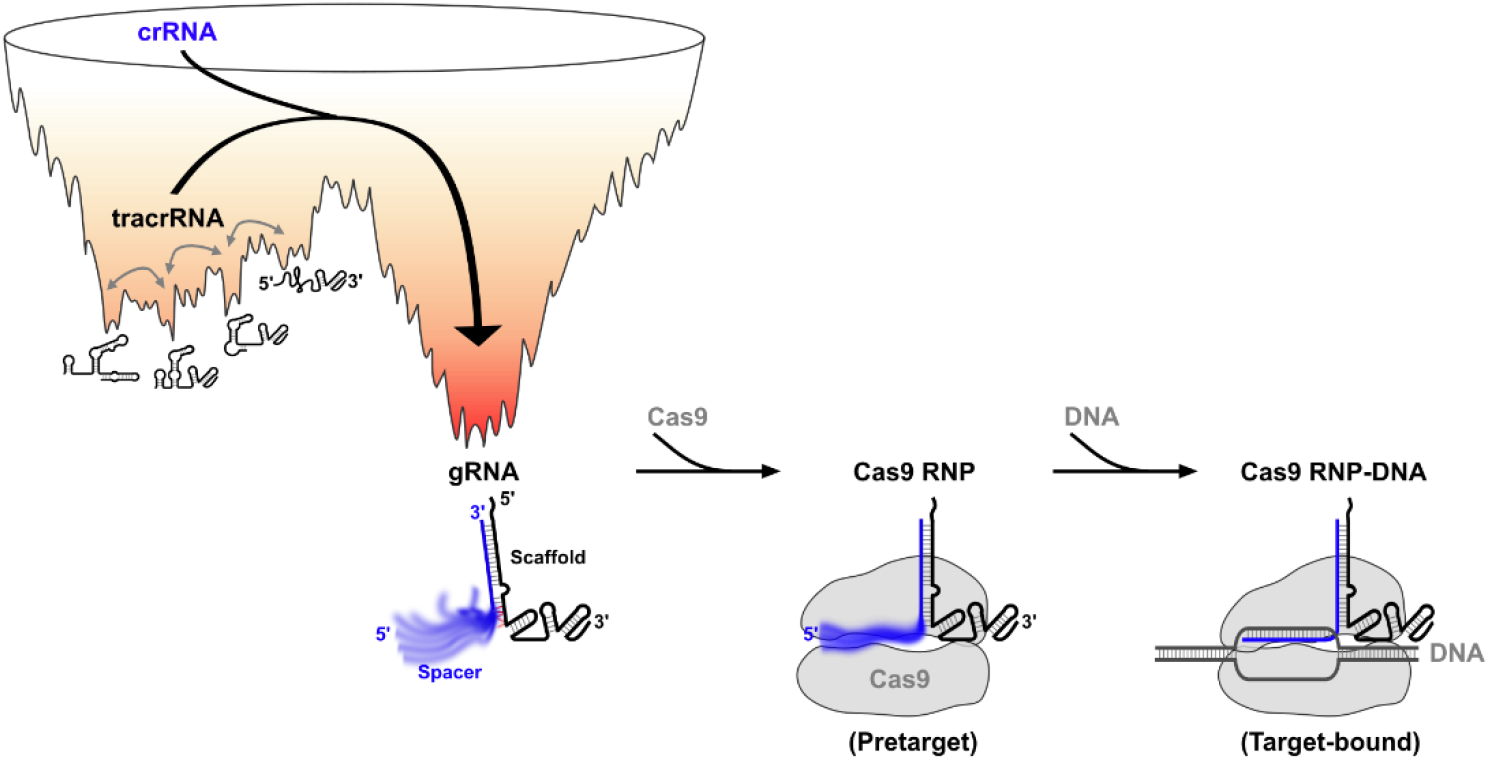
Model for gRNA initial folding and structural rearrangements along its assembly pathway. The transition from tracrRNA, which exists as an ensemble of interconverting structures to gRNA with crRNA, is thermodynamically favorable due to the stable partial duplex formation. The hybridized gRNA (*apo*) has a rigid RNA-RNA (scaffold) with a 5’ end flexible extension (spacer). In the pretarget state, the 5’ end spacer is still flexible. In the target-bound state, spacer transits to a rigid state as a result of RNA-DNA heteroduplex formation with its target DNA.

Our measurements revealed that the 5’ end tracrRNA where crRNA binds to form a partial duplex within the gRNA scaffold initially exists as an ensemble of interconverting conformations (Fig. 2A & B). This implies that the conformations of 5’ end tracrRNA ensemble are unstable and distribute on the surface of energy landscape at similar energy levels with relatively low energy barriers among them (Fig. 5). Once this 5’ end tracrRNA hybridized with the 3’ end crRNA, we observed one major static FRET population (Fig. 2A & B), which corresponded to the RNA-RNA duplex within the gRNA scaffold area. These observations suggest that the unstable ensemble of 5’ end tracrRNA can be occasionally unfolded and lead to folding into a thermodynamically stable gRNA partial duplex in the presence of crRNA (Fig. 5). Although we do not measure the 3’ end crRNA, we hypothesize that its conformation is also relatively unstable compared to when it is in the form of partial duplex of gRNA.

Unlike the stable duplex scaffold, the sequence variable spacer at the 5’ end gRNA was shown to be less stable and structurally flexible (Fig. 3D and S5). Therefore, ABEL-FRET measurements for the sequence-conserved scaffold (tracrRNA) and sequence-variable spacer (crRNA) regions depict the gRNA structure that has a flexible 5’ end spacer next to the well-defined duplex scaffold (Fig. 5). Furthermore, our analysis revealed that the 5’ end gRNA undergoes constant fluctuations at the sub-second timescales (∼ 0.3 second) in the low FRET state (0.25-0.35) with occasional transitions to mid-to-high FRET (0.6-0.8) states (Fig. 3D and S5). This suggests that the 5’ end spends most of its time in an extended form with brief transitions into more compact or random coil states (Fig. 5). The timescales observed for low FRET fluctuations fall in the range for RNA dynamics that are associated with localized base-paring fluctuations (microseconds to seconds) or larger-scale secondary structure transitions (> 0.1 second)^23^. Since we observed continuous fluctuations within small amplitude in the real-time FRET trace (Fig. 3D and S5), the 5’ end dynamic behaviors likely involved local changes in base pairs rather than secondary structure transitions which are usually shown as distinct FRET transitions. Perhaps, base pairs located at the basal region of the partial duplex stem, which also accounted for 5’ end FRET measurements based on our gRNA design (Fig. 3A), underwent constant breaking and reforming due to high local instability and caused slow motion at the extended 5’ end gRNA (Fig. 5). In contrast, the upper stem of partial duplex appeared to be relatively static and stable according to our measurements for tracrRNA when it was in a partial duplex gRNA (Fig. 2B, right trace). Therefore, our observations suggest that the long and highly base-paired partial duplex is critical for maintaining the overall structural integrity of gRNA for Cas9 recognition by forming a stable structural core that resists total denaturation, even when subjected to basal melting. As a result, gRNA in the *apo* state already adopts a pre-organized structure that resembles its Cas9-bound conformation as the stable partial duplex isolates the 5′ spacer from the 3′ scaffolding structure (Fig. 5).

Upon Cas9 binding, the *E*_FRET_ value of partial duplex was shifted from 0.42 to 0.49 (Fig. 2A), indicating the additional rearrangement at the gRNA scaffold by interactions with Cas9. In the crystal structure, this partial duplex has extensive interactions with Cas9^11,24^, indicating its crucial role in RNP complex formation^24,25^. On the other hand, the 5’ end gRNA spacer preserved its flexibility even after the Cas9 binding but with millisecond dynamics (Fig. 4C and D), which characteristically involve rapid and small-scale structural fluctuations within stable RNA structures (pico-to microseconds)^23^. Previous crystal and electron microscopy structures of Cas9-sgRNA indicated that the 20-nt spacer at the 5′ end gRNA is straightened along the positively charged cleft between two nuclease domains (HNH and RuvC) of Cas9^14^ with extensive contacts through its phosphate backbone^11^. As a result, the seed sequence (#11-20 nt) becomes less flexible in a preordered A-form like helical conformation, while the 5′ 10-nt nonseed sequence (#1-10 nt) is still disordered (Fig. 4A)^11^. Considering these previous structural works, our observations with ABEL-FRET for the pretarget-bound 5’ end spacer with highly uniformed millisecond dynamics reflect that the flexibility at the 5’ end gRNA was fine-tuned in the RNP complex. (Fig. 5). As a result of Cas9 binding, the 5’ end spacer was stabilized and captured in the functional shape for the following target DNA recognition but remained flexible within the limited Cas9-bound space, allowing it to adopt further conformational changes.

When the Cas9 RNP bound to target DNA, there was no significant effect on *E*_FRET_ of partial duplex scaffold, leading to indistinguishable *E*_FRET_ values between the pretarget- (0.49) and target-bound state (0.50) as shown in Fig. 2A. This implies the resembled configurations of gRNA at the partial duplex in the pretarget-and target-bound states, which is consistent with previous structural studies^24^. Although the 5’ end spacer stayed in the low FRET states for the entire assembly processes (Fig. 3C), we observed dramatic changes in its dynamic behaviors upon target binding that led to the conformational switching to the static RNA-DNA heteroduplex (Fig. 4B and C). Therefore, both scaffold and spacer of gRNA are structurally ordered in the target-bound state, giving rise to gRNA in the thermodynamically favored native structure (Fig. 5) that can further induce conformational rearrangements within Cas9 required for target cleavage.

Collectively, our ABEL-FRET visualizes the gRNA structures and dynamics from two viewpoints along its assembly into Cas9 RNP and reveals a dramatic transition from a highly flexible, structurally diverse, and rapidly rearranging ensemble to a stable, rigid, and functional complex. We anticipate that our approach can be applied to other RNP systems and therefore fill gaps in knowledge, particularly regarding how RNA structural dynamics and conformational changes influence the sequential bindings along the RNP assembly.

## Materials and Methods

### Cas9 and oligonucleotides

*S. pyogenes* Cas9 (#M0386T) and dCas9 (#M0652S or (#M0652T) were commercially purchased from New England Biolabs (NEB, Ipswich, MA) and used without further purifications. Both tracrRNA and sgRNA were synthesized by Bio-Synthesis (Lewisville, TX) with a 5’Cy3 and an amino modification at the designated uracil residue with a C6 linker (Amino C6U). The crRNA and target dsDNA were purchased with a 5’ Amino C12 (dsDNA) or without any modification (crRNA and target dsDNA) from Integrated DNA Technologies (IDT). The sequences of all strands used in this study are listed in Supplementary Table 1.

### Fluorescence labeling

Both 5’Cy3-attached tracrRNA and sgRNA were further labeled through the Amino C6U by incubating with ∼10-fold excess of sCy5 (Lumiprobe) in 0.2 M sodium bicarbonate buffer (pH 8.3) and 7 M urea overnight. The target DNA strands were labeled under nondenaturing conditions without urea. The labeled DNA and RNA strands were then purified twice using size-exclusion columns (BioRad P6). The labeling efficiencies were calculated using absorption measurements, which were determined to be 50–70% for RNA and 95-100 % for DNA.

### gRNA assembly

First, 1 μM of labeled tracrRNA without (tracrRNA only) or with 2 μM crRNA (crRNA+tracrRNA) was heated at 94 °C for 5 minutes, followed by slowly cooling to 25 °C at - 1°C/min rate in 20 mM HEPES (pH 8) with 100 mM KCl. The labeled sgRNA (1 μM) was also refolded using the same temperature cycle. The Cas9 RNP formation was induced by incubating 100 nM of gRNA (crRNA+tracrRNA or sgRNA) and 200 nM of protein (Cas9 or dCas9) at 25 °C for 10 minutes in 1X NEBuffer™ r3.1 (#B7203S) that contained 100 mM NaCl, 50 mM Tris-HCl, 10 mM MgCl_2_, 100 µg/mL recombinant albumin (pH 7.9). For the target-bound RNP formation, this mixture was further incubated at 37°C for 15 minutes with 1 μM of pre-annealed target dsDNA.

### DNA cleavage assays

The 5’sCy3-attached DNA (target and nontarget) strands were annealed by thermal cycles, which gave dsDNA labeled at the 5’ ends of both target and nontarget strands. The preassembled RNP complexes (Cas9 or dCas9 RNP) with equivalent amount of gRNA and enzyme at 100 nM (25 °C, 10 min) were incubated with 10 nM of labeled dsDNA in NEBuffer™ r3.1 at 37°C for 15 minutes. The reactions were stopped by adding 0.5 µL of RNase A (10 mg/mL, ThermoFisher Scientific, EN0531) and 0.5 µL of protease K (800 units/mL, NEB, #P8107S). The products were separated using native polyacrylamide gel electrophoresis (native-PAGE) and visualized for sCy5 using Typhoon (Amersham).

### Sample preparation for ABLE-FRET experiments

Pre-mixed samples were diluted to have a final concentration of 5 pM labeled RNA (tracrRNA or sgRNA). All measurements were performed in a buffer containing 20 mM HEPES (pH 8), 100 mM KCl, 5 mM MgCl_2_, and 3 mM Trolox in the presence of enzymatic oxygen scavenger system with 2.5 mM protocatechuic acid (Sigma) and 50 nM protocatechuate-3,4-dioxygenase (OYC America, purified by size exclusion chromatography).

### Surface preparation

The microfluidic devices^26,27^ were cleaned multiple times in piranha solution (3:1 mixture of sulfuric acid and hydrogen peroxide), with extensive rinsing in ultrapure water (18.2 MΩ) after each cleaning cycle. To prevent nonspecific adsorption of protein, device surfaces were coated with poly(ethylene glycol) (PEG) by first incubating in 1 M potassium hydroxide for 15 min, followed by incubation in mPEG–silane solution (Laysan Bio MPEG-SIL-5000–1g, >20 mg·ml^-1^ in 95% ethanol–5% water mixture with pH ∼5)^28^ for more than 24 hours at room temperature. PEGylated surfaces were then rinsed extensively with ultrapure water and dried with pure nitrogen before use.

### ABEL-FRET setup

ABEL-FRET, which integrates an ABEL trap^5^ with photon-counting smFRET detection optics, was implemented based on a previously published design^29^. In this setup, single molecules freely diffusing within the trapping region (∼3×3 μm^2^) of the microfluidic device are monitored photon-by-photon. Molecular position is estimated in real-time via rapid beam scanning and a Kalman filter. The resulting voltage feedback is amplified and applied instantaneously to counter Brownian motion, enabling the molecule to remain trapped for extended time (up to 15 s in this study). While trapped, photons emitted from the molecule are collected by the confocal microscope, which separated accordingly into donor and acceptor detection channels, and time-stamped for smFRET measurements. Once the donor fluorophore photobleaches, the system releases the molecules and resets to capture the next one.

### Hydrodynamic profiling

Diffusion coefficient (*D*) and electrokinetic mobility (*µ*) of isolated single molecules were extracted from the position estimates and voltage feedback generated during ABEL trapping, as previously described^31^. Briefly, photon-stamped position estimates and corresponding feedback voltages are segmented into independent 100-200 ms blocks. An expectation-maximization algorithm, operating with a maximum likelihood framework, processed these positions and voltages data to reconstruct the in-trap motion trajectory of each single molecule. This reconstructed trajectory was then decomposed into a voltage-dependent component and a stochastic component, which *µ* and *D* were estimated, respectively.

### FRET efficiency calculation

Time-tagged photon-by-photon data were first converted into “mcs” traces by binning into fixed time intervals (counts/bin time), yielding traces that contain both single-molecules signal and background regions. The start and end time points of each signal and background region were identified using the change-point finding algorithm^32,33^. Background photon-count histograms from background regions of each trace were fitted with a Poisson distribution to determine the mean background rates for the donor, acceptor, and combined channels. Single-molecules signal regions included molecules exhibiting active FRET pairs, donor-only (no acceptor), and acceptor-photobleached populations. Background-corrected photon counts from donor (*N*_D_) and acceptor channels (*N*_A_) were used to calculate the apparent *E*_FRET_ of single molecules, incorporating donor leakage (*α*) and detection (*γ*) correction factors, using the following equations:

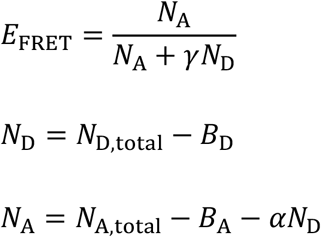

where *N*_D, total_ and *B*_D_ are the total and background photon counts (5 ms bin time) on the donor channel, *N*_A, total_ and *B*_A_ are the counts on the acceptor channel, and *α* and *γ* are the correction factors described below. The *α* factor was determined by fitting the donor-only population with a Gaussian distribution to have the mean *E*_FRET_ of 0. The *γ* factor was determined by Gaussian fitting of the intensity (counts/5ms) histograms for donor-only and FRET populations to have the equivalent mean values of intensity for both populations using the following equation:

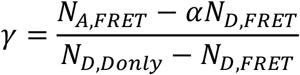

where *N*_D, FRET_ and *N*_A, FRET_ are the mean photon counts for the FRET population from the donor and acceptor channels, and *N*_D, Donly_ is the mean photon counts for the donor-only population from the donor channel.

### Cross-correlation analysis

Intensity cross-correlation between the donor and acceptor channels was calculated from the time-tagged photon-by-photon data, binned at 10 ms for free sgRNA (*apo*) or at 2 ms for Cas9-bound sgRNA (pretarget). The resulting cross-correlation curves were fitted with a single exponential function^34^ with an offset:

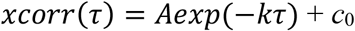

where *A* and *k* are the two fitting parameters, *τ* is the time lag over which the correlation is calculated, and *c*_0_ is a floating offset term. Time lag intervals were chosen to align with the beam scanning rate to avoid periodic artifacts caused by the physical movement of the laser beam.

### FRET change point analysis

Change points in single-molecule FRET traces of 5’ end-labeled sgRNA in the *apo* state were identified using a previously developed change point algorithm^35^. Briefly, the baseline noise of each FRET trace was first shifted to a mean of zero. Change points were then identified using a binary segmentation algorithm under a Gaussian noise model, with a fixed noise variance (σ^2^ = 0.0015) and a significance threshold (α = 0.05) applied to the log-likelihood ratio test statistic. Between consecutive change points, discrete FRET states were defined as the mean FRET value within each segment.

## Notes

### Competing Interest Statement

The authors have declared no competing interest.

## Reference

(1) Goodrich, J. A. Non-Coding-RNA Regulators of RNA Polymerase II Transcription. Nat. Rev. Mol. Cell Biol. 2006, 7, 612–616.

(2) Cech, T. R. The Noncoding RNA Revolution—Trashing Old Rules to Forge New Ones. Cell 2014, 157 (1), 77–94. 10.1016/j.cell.2014.03.008.

(3) Chauvier, A. Monitoring RNA Dynamics in Native Transcriptional Complexes. Proc. Natl. Acad. Sci. 2021, 118 (45).

(4) Xue, Y. Observation of Structural Switch in Nascent SAM-VI Riboswitch during Transcription at Single-Nucleotide and Single-Molecule Resolution. Nat. Commun. 2023, 14, 2320.

(5) Liao, T.-W. Linking Folding Dynamics and Function of SAM/SAH Riboswitches at the Single Molecule Level. Nucleic Acids Res. 2023, 51 (17), 8957–8969. 10.1093/nar/gkad633.

(6) Wilson, H.; Wang, Q. ABEL-FRET: Tether-Free Single-Molecule FRET with Hydrodynamic Profiling. Nat. Methods 2021, 18 (7), 816–820. 10.1038/s41592-021-01173-9.

(7) Cohen, A. E. Method for Trapping and Manipulating Nanoscale Objects in Solution. Appl. Phys. Lett. 2005, 86 (9), 093109. 10.1063/1.1872220.

(8) Wang, Q. Probing Single Biomolecules in Solution Using the Anti-Brownian Electrokinetic (ABEL) Trap. Acc. Chem. Res. 2012, 45 (11), 1955–1964.

(9) Jinek, M.; Chylinski, K.; Fonfara, I.; Hauer, M.; Doudna, J. A.; Charpentier, E. A Programmable Dual-RNA–Guided DNA Endonuclease in Adaptive Bacterial Immunity. Science 2012, 337 (6096), 816–821. 10.1126/science.1225829.

(10) Jinek, M. RNA-Programmed Genome Editing in Human Cells. eLife 2013, 2 (e00471).

(11) Jiang, F.; Zhou, K.; Ma, L.; Gressel, S.; Doudna, J. A. A Cas9–Guide RNA Complex Preorganized for Target DNA Recognition. Science 2015, 348 (6242), 1477–1481. 10.1126/science.aab1452.

(12) Jinek, M. Structures of Cas9 Endonucleases Reveal RNA-Mediated Conformational Activation. Science 2014, 343 (6176). 10.1126/science.1247997.

(13) Nishimasu, H.; Ran, F. A.; Hsu, P. D.; Konermann, S.; Shehata, S. I.; Dohmae, N.; Ishitani, R.; Zhang, F.; Nureki, O. Crystal Structure of Cas9 in Complex with Guide RNA and Target DNA. Cell 2014, 156 (5), 935–949. 10.1016/j.cell.2014.02.001.

(14) Jiang, F.; Taylor, D. W.; Chen, J. S.; Kornfeld, J. E.; Zhou, K.; Thompson, A. J.; Nogales, E.; Doudna, J. A. Structures of a CRISPR-Cas9 R-Loop Complex Primed for DNA Cleavage. Science 2016, 351 (6275), 867–871. 10.1126/science.aad8282.

(15) Singh, D.; Sternberg, S. H.; Fei, J.; Doudna, J. A.; Ha, T. Real-Time Observation of DNA Recognition and Rejection by the RNA-Guided Endonuclease Cas9. Nat. Commun. 2016, 7 (1), 12778. 10.1038/ncomms12778.

(16) Dagdas, Y. S.; Chen, J. S.; Sternberg, S. H.; Doudna, J. A.; Yildiz, A. A Conformational Checkpoint between DNA Binding and Cleavage by CRISPR-Cas9. Sci. Adv. 2017, 3 (8), eaao0027. 10.1126/sciadv.aao0027.

(17) Singh, D. Mechanisms of Improved Specificity of Engineered Cas9s Revealed by Single Molecule FRET Analysis. Nat. Struct. Mol. Biol. 2018, 25 (4), :347-354.

(18) Wang, Y. Real-Time Observation of Cas9 Postcatalytic Domain Motions. Proc. Natl. Acad. Sci. 2020, 118 (2), e2010650118. 10.1073/pnas.2010650118.

(19) Okafor, I. C. Single Molecule FRET Analysis of CRISPR Cas9 Single Guide RNA Folding Dynamics. J. Phys. Chem. B 2022, 127 (1).

(20) Talkington, M. W. T. An Assembly Landscape for the 30S Ribosomal Subunit. Nature 2005, 438, 628–632.

(21) Stone, M. D. Stepwise Protein-Mediated RNA Folding Directs Assembly of Telomerase Ribonucleoprotein. Nature 2007, 446, 458–461.

(22) Doudna, J. A. The New Frontier of Genome Engineering with CRISPR-Cas9. Science 2014, 346 (6213). DOI: 10.1126/science.1258096.

(23) Mustoe, A. M. Hierarchy of RNA Functional Dynamics. Annu. Rev. Biochem. 2014, 83, 441–466. 10.1146/annurev-biochem-060713-035524.

(24) Jiang, F. CRISPR–Cas9 Structures and Mechanisms. Annu. Rev. Biophys. 2017, 46, 505– 529. 10.1146/annurev-biophys-062215-010822.

(25) Briner, A. E. Guide RNA Functional Modules Direct Cas9 Activity and Orthogonality. Mol. Cell 2014, 56 (2), 333–339.

(26) Cohen, A. E. Controlling Brownian Motion of Single Protein Molecules and Single Fluorophores in Aqueous Buffer. Opt. Express 2008, 16 (10), 6941.

(27) Manger, L. H. Revealing Conformational Variants of Solution-Phase Intrinsically Disordered Tau Protein at the Single-Molecule Level. Angew. Chem. Int. Ed. 2017, 56 (49), 15584–15588.

(28) Witucki, G. L. A Silane Primer: Chemistry and Applications of AIkoxy Silanes. J. Coat. Technol. 1993, 65, 57–60.

(29) Wilson, H. ABEL-FRET: Tether-Free Single-Molecule FRET with Hydrodynamic Profiling. Nat. Methods 2021, 18, 816–820.

(30) Wang, Q. An Adaptive Anti-Brownian Electrokinetic Trap with Real-Time Information on Single-Molecule Diffusivity and Mobility. ACS Nano 2011, 5, 5792–5799.

(31) Wang, Q. Single-Molecule Motions Enable Direct Visualization of Biomolecular Interactions in Solution. Nat. Methods 2014, 11, 555–558.

(32) Watkins, L. P. Detection of Intensity Change Points in Time-Resolved Single-Molecule Measurements. J. Phys. Chem. B 2005, 109 (1), 617–628.

(33) Wang, Q. Lifetime and Spectrally Resolved Characterization of the Photodynamics of Single Fluorophores in Solution Using the Anti-Brownian Electrokinetic Trap. J. Phys. Chem. B 2012, 117 (16), 4641–4648.

(34) Kim, H. D. Mg2+-Dependent Conformational Change of RNA Studied by Fluorescence Correlation and FRET on Immobilized Single Molecules. Proc Natl Acad Sci U A 2002, 99 (7), 4284–4289.

(35) Hugh, W. Joint Detection of Change Points in Multichannel Single-Molecule Measurements. J. Phys. Chem. B 2021, 125 (49).

